# Dorso-Ventral and Night-Day Regulation of Extracellular K+ Dynamics in Mouse Hippocampal Astrocytes

**DOI:** 10.1101/2025.10.10.681546

**Authors:** Nariman Kiani, Kabeer Abubakar, Tin Luka Petanjek, Monique Esclapez, Anton Ivanov, Christophe Bernard

**Affiliations:** Aix Marseille Univ, INSERM, INS, Inst Neurosci Syst, Marseille, France; University of Zagreb, Department of Physiology and Department of Neuroscience, Croatian Institute for Brain Research, School of Medicine, Zagreb, Croatia; Aix Marseille Univ, Inserm, INMED, Institut de Neurobiologie de la Méditerranée, Marseille, France

**Keywords:** Astrocytes, hippocampal regions, K+ homeostasis, Kir4.1 channels, circadian rhythm

## Abstract

Astrocytes regulate extracellular potassium (K^+^) through multiple mechanisms operating across distinct spatiotemporal scales, yet whether this regulation exhibits regional and circadian specificity remains unclear. Using real-time K^+^ measurements combined with electrophysiology and immunohistochemistry in astrocytes, we characterize K^+^ buffering by astrocytes across the dorsoventral hippocampal axis at three circadian times. We report that hippocampal CA1 *stratum radiatum* astrocytes possess different functional K^+^ buffering capacities depending on both anatomical location and time of day. At the early light phase (ZT3), the ventral hippocampus (VH) exhibits faster K^+^ accumulation and greater peak amplitudes than the dorsal hippocampus (DH), despite similar neuronal network activity. This regional divergence is driven by reduced Kir4.1 channel function in VH astrocytes, which persists across all measured time points, and which is partially compensated by enhanced Na^+^/K^+^-ATPase activity specifically at ZT3. Gap junction coupling between astrocytes shows regional and time-dependent variation, with elevated coupling in VH at ZT3 that subsequently normalizes. Kir4.1 protein expression exhibits circadian dynamics, particularly in DH where expression declines from early light to late light and recovers in early dark. These findings establish astrocytic K^+^ buffering as a multiscale phenomenon integrating regional heterogeneity with circadian regulation, with implications for understanding regional and temporal differences in network excitability, particularly in epilepsy.

## Introduction

Under physiological conditions, extracellular K^+^ concentration ([K^+^]ₒ) fluctuates between 2 and 5 mM, while increases above 10 mM are linked to pathological activities such as seizures and spreading depression (de Curtis et al., 2018; Enger et al., 2015; Erecińska & Silver, 1994). Maintaining [K^+^]ₒ within narrow physiological limits is therefore functionally important.

Astrocytes play an important role in regulating [K^+^]ₒ by taking up excess extracellular K^+^, redistributing it through gap-junction–connected networks, and extruding it where [K^+^]ₒ is lower (Wallraff et al., 2006; Walz, 2000). This process relies on inward-rectifying Kir4.1 channels (Seifert et al., 2009), and the Na^+^/K^+^-ATPase pump. Impairment of either component can increase neuronal and network activity and excitability (Djukic et al., 2007; Romanos et al., 2020; Wallraff et al., 2006).

Neuronal activity and excitability are not uniform across the brain. Even within a single structure, distinct subregions exhibit marked differences in intrinsic excitability, synaptic properties, and responses to pharmacological or pathological challenges. For example, along the dorsoventral axis of the hippocampus, the ventral region (VH) displays higher intrinsic excitability, greater seizure susceptibility, distinct patterns of synaptic plasticity, and different metabolic properties compared to the dorsal region (DH) (Brancati et al., 2021; Dougherty et al., 2012; Papatheodoropoulos et al., 2005). Similarly, neuronal activity is not constant across time: circadian rhythms modulate neuronal excitability, synaptic transmission, and seizure threshold (B.-L. Chang et al., 2018; Debski et al., 2020; Quigg et al., 2000), reflecting temporal regulation of molecular, metabolic, and physiological processes (Brancati et al., 2021; Debski et al., 2020; McCauley et al., 2020).

This raises a fundamental question: do astrocytes exhibit similar spatial and temporal heterogeneity in their K^+^ buffering capacity? Do they "follow the pace" of neuronal activity, adapting their homeostatic functions to match regional differences in excitability and circadian variations in network state? If astrocytic K^+^ buffering were spatially and temporally invariant, regions of high neuronal excitability (such as the VH) and periods of elevated seizure susceptibility (such as specific circadian phases) would be vulnerable to inadequate K^+^ homeostasis. Conversely, if astrocytes dynamically adjust their buffering capacity in space and time, this could represent a sophisticated form of adaptive homeostasis matched to local circuit demands.

The hippocampus provides an ideal system to address this question. Beyond the well-documented differences in neuronal properties along the dorsoventral axis and across circadian time, the hippocampus also exhibits regional and temporal heterogeneity in astrocytic molecular markers, morphology, and calcium signaling (Dong et al., 2009; Jinno, 2011; Ryu et al., 2024; Thompson et al., 2008). However, whether these molecular and morphological differences translate into functional differences in K^+^ buffering—and how they relate to the known regional and circadian variation in neuronal excitability—remains unclear.

Here, we directly test whether astrocytic K^+^ buffering capacity varies across the dorsoventral hippocampal axis and across circadian time, using integrated functional, electrophysiological, and molecular approaches. We use an extracellular K^+^ sensor to measure K^+^ dynamics during standardized synaptic stimulation (Schaffer collateral activation), monitoring multiple parameters of the K^+^ transient (accumulation rate, peak amplitude, clearance kinetics, and Na^+^/K^+^-ATPase-dependent undershoot) in dorsal and ventral hippocampal slices at three Zeitgeber times: ZT3 (early light, baseline period), ZT8 (late light, period of heightened seizure susceptibility (W.-C. Chang et al., 2018; Debski et al., 2020; Quigg et al., 2000), and ZT15 (early dark, active phase). We then employ whole-cell patch-clamp electrophysiology to assess the cellular mechanisms underlying these functional differences, examining gap-junction coupling, Kir4.1 channel function, and astrocyte resting membrane potential. Finally, we use immunohistochemistry to quantify regional and circadian variation in Kir4.1 protein expression. Our findings demonstrate that astrocytic K^+^ buffering capacity depends upon both anatomical location and circadian phase, revealing a form of adaptive heterogeneity that parallels—but does not simply mirror—the spatial and temporal organization of neuronal excitability.

## Methods

### Animals and Ethical Approval

All animal procedures were conducted in accordance with the European Directive 2010/63/EU and French regulations on animal experimentation. The study protocol was approved by the relevant institutional and national ethics committees (APAFIS authorization #44598-2023081812555435v4). A total of 50 male FVB mice (weight: 28–45 g; age: 2 months or older) were used. Of these, 3 mice were dedicated to immunohistochemistry, while the remaining animals were used for both patch-clamp experiments and K^+^ dynamic recordings in parallel. The single-sex design increased statistical power and ensured an adequate yield of viable samples given the technically demanding nature of the experiments. The exclusion of female mice is a limitation of the study, and future work will be required to assess the generalizability of these findings across sexes.

### Brain slice preparation

Mice anesthetized with sevoflurane, were decapitated, and the brain was swiftly removed from the skull and submerged in ice-cold artificial cerebrospinal fluid (ACSF) (in mM): NaCl 126, KCl 3.50, NaH_2_PO_4_ 1.25, NaHCO_3_ 25, CaCl_2_ 2, MgSO_4_1.3, and glucose 10, with a pH value maintained between 7.3 and 7.4. The ACSF was aerated with a 95% O2/5% CO2 gas mixture. One hemisphere, left or right in alternation, was used to prepare dorsal slices via coronal sections, whereas the other hemisphere was used to prepare ventral slices as described previously (Dougherty et al., 2012). The brain was immersed in an ice-cold cutting solution (in mM): K-gluconate 140, HEPES 10, Na-gluconate 15, EGTA 0.2, and NaCl 4, with the pH adjusted to 7.2 with KOH. Using a tissue slicer (Leica VT 1200s, Leica Microsystem, Germany), 300 μm-thick for patch-clamp recording and 350 μm- thick slices for field recording were produced. Subsequently, to preserve the cells near the surface from resulting damage caused by slicing, the 300 μm-thick slices were incubated for 15 to 20 minutes at room temperature (RT) in a choline chloride solution (in mM): Choline Chloride 110, KCl 2.5, NaH_2_PO_4_ 1.25, MgCl_2_ 10, CaCl_2_ 0.5, NaHCO_3_ 25, Glucose 10, Na-Pyruvate 5, constantly bubbled with a 95% O_2_/5% CO_2_ gas mixture. To label the astrocytes, the slices were incubated for 20 minutes in a ACSF solution containing 1 µM of sulforhodamine 101 (SR101, Merk), a fluorescent dye specifically absorbed by astrocytes and not neurons (Nimmerjahn et al., 2004). The slices were allowed to recover in a dual-side perfusion holding chamber with continuously circulating ACSF for at least one hour before experimentation.

### Synaptic stimulation and field potential recordings

In parallel, 350 μm-thick transferred to a dual perfusion recording chamber continuously perfused (∼6 ml/min) with ACSF containing 5 mM of glucose, warmed to 30°C. Schaffer collateral (SC) was stimulated with a bipolar metal electrode connected to a DS2A-isolated stimulator (Digitimer Ltd, UK). The electrode tip was placed in the *Stratum Radiatum* (*s.r.*) between the CA3 and CA2 regions. Extracellular recording electrodes were pulled (Harvard Apparatus Ltd, UK, O.D. = 1.2 and I.D. = 0.94 mm, Rp < 1 MΩ) using a horizontal puller (Sutter Instruments, USA). The extracellular local field potential (LFP) was recorded with a glass microelectrode filled with ACSF, placed in the *Stratum Oriens* (*s.o.*) towards the *Stratum Pyramidale* (*s.p.*) of the CA1 area. The signal was amplified 1000 times with an EXT-02F amplifier (NPI Electronic, Germany) operating with a low-pass 3kHz filter (DC mode). The stimulation current intensity was adjusted to produce an LFP response of 60-70% of the maximum amplitude. Each slice was stimulated with trains of stimulation pulses of different frequencies ranging from 2.5Hz to 100Hz. The LFP responses were recorded simultaneously with the K^+^-dependent potential using Patch Master software. Each stimulation protocol was repeated twice. The LFP responses to single pulse stimulation were monitored in between train stimulations to check the slice stability. If the response decreased by more than 20%, the slice was discarded. In this study, we used the response to the 10Hz30s trains because this stimulation evoked a K^+^ signal with good SNR, allowing the measurement of required parameters (peak amplitude, 25-75% rise time, decay time, and undershoot amplitude) of the individual and not the averaged response.

### Fabrication of Potassium Sensitive Microelectrode

K+-selective microelectrodes were prepared using the method described by Heinemann and Arens (1992). In brief, electrodes were pulled from double-barrel theta glass (TG150-4, Warner Instruments, Hamden, CT, USA). The reference barrel was filled with 154 mmol/L NaCl solution. The silanized ion-sensitive barrel tip (5% trimethyl-1-chlorosilane in dichloromethane) was filled with a potassium ionophore I cocktail A (60031 Fluka distributed by Sigma-Aldrich, Lyon, France) and backfilled with 100 mmol/L KCl. Measurements of K^+^-dependent potentials were performed using a high-impedance differential DC amplifier equipped with negative capacitance feedback control, which permitted the compensation of the microelectrode capacitance required for the recording of the rapid changes of [K^+^]ₒ. The electrodes were calibrated before each experiment. The calibration curve, concentration vs potential, was fitted with an exponential function. The obtained equation was used to convert [K^+^]ₒ-dependent potential recordings into concentration changes.

### Electrophysiological recording of the astrocytes

For the electrophysiological recordings of astrocytes, 300 μm-thick slices were immersed in a low-volume (2 ml) recording chamber and continuously perfused with ACSF at 32 - 34°C and a perfusion rate of 5 mL/min. Sulforhodamine labeled astrocytes in the *stratum radiatum* (*s.r.*) of the CA1 region of the hippocampus were identified as a fluorescent cells with astrocytic morphology (excitation/emission wavelengths of 586/605 nm). Patch pipettes were fabricated using the same horizontal puller from borosilicate glass tubing (Harvard Apparatus Ltd, UK,1.5 mm outer diameter (O.D.), 0.86 mm inner diameter (I.D.), pipette resistance (Rp) = 5.78 ± 0.12 MΩ) and filled with an intrapipette solution (in mM): KCl 20, K-gluconate 115, HEPES 10, EGTA 1.1, MgATP 4, Na -phosphocreatine 10, Na_2_GTP 0.4, and 0.3-0.4% biocytin (Sigma, Germany). Signals were transmitted to a Multiclamp 700A amplifier (Molecular Devices), digitized at a rate of 10 kHz using a DigiData 1550 interface (Molecular Devices) connected to a personal computer, and analyzed using ClampFit software (Molecular Devices). The resting membrane potential (RMP) of astrocytes was measured using the patch-clamp technique, whole-cell configuration, in current clamp mode.

### Isolation of Kir4.1 currents

To isolate Kir4.1 currents and to keep constant the extracellular K^+^, slices were incubated for 36 ± 2 minutes in the baseline 5 mM glucose-containing ACSF supplemented with (in µM): Meclofenamic acid (MFA, Sigma-Aldrich) 100 to block gap junctions, and Tetrodotoxin (TTX, Abcam) 0.5, 6-Cyano-7-nitroquinoxaline-2,3-dione (CNQX, HelloBio) 10, and D-(-)-2-Amino-5-phosphonopentanoic acid (D-AP5, HelloBio) 40 to stop neuronal firing and minimize synaptic transmission. No effect of incubation time on Kir4.1 current was detected. Upon establishment of the whole-cell patch clamp configuration, the patched astrocyte was held at -70 mV in voltage clamp mode, and after 5 minutes of dialysis, membrane currents were recorded at different holding potentials varying from -140mV to 50mV with 10mV steps. Then, to block Kir4.1 channels, 100 μM of BaCl_2_ was added to the baseline ACSF, and the currents elicited by the same voltage steps were recorded after 5 minutes. The current-voltage relationship of Ba^2+^ sensitive conductance was determined by subtracting the currents recorded in Ba^2+^ from those recorded before its administration. Access resistance was determined by biexponentially fitting the capacitive current resulting from −70 to −60 mV voltage step. Cells were discarded if they met any of the following criteria: an initial access resistance above 35 MΩ, a resting membrane potential (RMP) less negative than -50 mV, or a final access resistance increase of more than 20%.

### Coupling estimation

Astrocytes are extensively coupled via gap junctions formed by Cx43 and Cx30, supporting intercellular redistribution of ions and metabolites, including potassium buffering. Pharmacological inhibition of these gap junctions is expected to uncouple astrocytes and thereby reducing their capacity for spatial potassium buffering. In principle, comparing K^+^-dependent currents evoked by Schaffer collateral stimulation before and after gap junction blockade could provide an estimate of the coupling state of the recorded astrocyte.

However, available gap junction inhibitors lack specificity and profoundly affect neuronal and glial function. Carbenoxolone alters both AMPA and GABA_A_ receptor-mediated synaptic transmission (Tovar et al., 2009). Meclofenamic acid is a potent activator of KCNQ2/Q3 (Kv7.2/7.3, M-type) channels (U. A. Anderson et al., 2009). and, at the 50–100 µM concentrations commonly used to block gap junctions, hyperpolarizes CA1 neurons and suppresses synaptic responses (Peretz et al., 2005). Connexin mimetic peptides such as

Gap26 and Gap27, while they are more specific at the channel level, also reduce basal excitatory synaptic transmission, likely by inhibiting gliotransmitter release through Cx43 hemichannels (Dospinescu et al., 2025). Because all of these agents can alter overall network activity, they inevitably affect not only K+ buffering but also activity-dependent K+ release. To avoid these confounding effects, we adopted an alternative approach to assess astrocytic coupling, based on the spread of the gap junction–permeable intracellular tracer biocytin following controlled whole-cell loading (Stephan et al., 2021).

In a separate set of experiments, astrocytes were patched in the absence of any blocker in whole cell configuration and kept at -70 mV holding potential for 20 mins. Slices were then fixed in 4% paraformaldehyde (AntigenFix, Diapath) overnight at 4°C. After three 30-minute washes in 0.12 M phosphate buffer (PB), slices were cryoprotected in 20% sucrose overnight, then deep-frozen on dry ice and stored at -80°C until the day of experimentation.

To reveal biocytin-containing astrocytes, slices were rinsed in PB twice, in 0.02 M potassium phosphate buffered saline (KPBS) three times for 30 min, and then incubated with streptavidin-conjugated with fluorophore Alexa 488 (Abcam) diluted (1/200) in KPBS containing 0.3% Triton-X100, overnight at RT. After washing with KPBS for 3 x 30 min, slices were mounted on Superfrost™ slides using Fluoromount-G.

To visualize the stained astrocytes coupling, slices were imaged using a Zeiss LSM 700 inverted confocal microscope with a 20x (0.5x zoom) objective. Z-stacks (15–35 optical slices, length: 2048 pixels, width: 2048; 1 µm interval) were acquired in the CA1 *stratum radiatum* using Zeiss ZEN software. Z-stacks were merged using Fiji software (NIH), and pixel intensity was then normalized.

A blinded examiner then counted the number of stained astrocytes. Since only one astrocyte per slice was patched, the stained astrocytes were mandatory, those connected with the patched one directly or through other connected astrocytes. Therefore, we considered the number of stained astrocytes as an estimation of the astrocytic coupling.

### Immunohistochemistry and image analysis

#### Tissue preparation, Immunohistochemistry, and image analysis

Wild-type male FVB mice (P50–70) were anesthetized with an intraperitoneal injection of ketamine and xylazine, followed by a lethal dose of pentobarbital administered at designated Zeitgeber times (ZT3, ZT8, or ZT15). Mice were then transcardially perfused with cold 0.12 M PB to remove the blood and then with AntigenFix to fix the tissue. Brains were extracted, post-fixed for up to 4 hours in the same fixative, rinsed in PB, cryoprotected in 20% sucrose in PB overnight at 4°C, quickly frozen on dry ice and kept at -80°C. Hemisected blocks of brain containing dorsal and ventral hippocampus were sectioned at 40 µm using a cryostat. The Sections were collected sequentially in tubes containing an ethylene glycol-based cryoprotective solution (Lu & Haber, 1992; Watson et al., 1986) and stored at –20°C until histological processing.

#### Double labeling for GFAP and Kir4.1

Sections were processed for the simultaneous detection of GFAP to label astrocytes and the Kir4.1 channel. They were rinsed for 30 min in PB and for 2x30 mins in KPBS. Sections were then incubated for 1h at RT in 3% normal donkey serum (NDS) diluted in KPBS containing 0.3% Triton X-100 and overnight at RT in a solution containing the polyclonal guinea pig anti -GFAP (1:600, #AFP-001-GP, Alomone Labs) and polyclonal rabbit anti-Kir4.1 (1:500, #APC-035, Alomone Labs) diluted in KPBS containing 0.3% Triton X-100 and 1% NDS. After 3, 30-minute rinses in KPBS, sections were incubated for 2 h in Alexa Fluor 488-conjugated donkey anti-guinea pig IgG (1:100, #706-545-148, Jackson ImmunoResearch) and Cy3-conjugated donkey anti-rabbit IgG (1:100, #711-167-003, Jackson ImmunoResearch), diluted in KPBS and 3% NDS. Sections were then rinsed with 3x30 rinses of KPBS and mounted on Superfrost™ slides using Fluoromount-G containing the nuclear marker DAPI (Thermo Fisher).

Immunofluorescence was imaged on a Zeiss LSM 700 inverted confocal microscope using a 63x oil-immersion objective. Z-stacks (12–19 optical slices at 1 µm intervals) of the CA1 *stratum radiatum* were acquired using Zeiss ZEN software. One field of view per section was imaged. Z-projections were performed in Fiji (NIH), and GFAP signal was quantified by measuring its integrated density. Thereafter, the GFAP was thresholded using triangle method, creating a mask that defined the region of interest (ROI) for Kir4.1 analysis. Then, Kir4.1 signal within the GFAP-defined ROI was quantified using the AND operation in the Image Calculator, and the Kir4.1/GFAP integrated density ratio was calculated (Bataveljic et al., 2024).

### Statistical analysis

Statistical analyses were performed using R software (version 4.5.1). K^+^ transient parameters were all normalized to that of DH at ZT3, and subsequent analyses were performed on these normalized data. To provide a comprehensive analysis of the data, we used two different approaches: permutation-based hypothesis testing to identify significant main effects and Bayesian multilevel modeling to estimate the magnitude and uncertainty of those effects.

#### Permutation-Based Hypothesis Testing

Analyses involving the effects of region (DH vs. VH) and time (ZT3, ZT8, and ZT15) were performed using a resampling-based, Freedman-Lane permutation ANOVA from the permuco package in R (Kherad-Pajouh & Renaud, 2015). This non-parametric approach was selected to provide robust inference by reducing assumptions about data distribution—such as normality and sphericity. The analysis was conducted with 10,000 permutations to ensure stable p-value estimates. Effect sizes were calculated and reported as omega squared (ω²), which represents the proportion of variance in the tested parameter accounted for by each effect in the population. The magnitude of regional differences was also evaluated using an online estimation statistics tool, which reported the mean difference and corresponding 95% confidence intervals (Ho et al., 2019).

#### Bayesian Multilevel Modeling

To further analyze the effects of region, time, and their interactions while accounting for the hierarchical structure of our experimental design (e.g., multiple cells nested within animals), we performed Bayesian multilevel modeling using the brms package (version 2.22.0) (Bürkner, 2017). This framework allowed us to move beyond binary significance testing to provide a probabilistic estimation of parameter uncertainty. Models were specified with fixed effects for region, timepoint, and their interaction.

We used the fitdistrplus package (version 1.2.4) to identify the optimal family distribution for each variable based on the lowest Akaike Information Criterion (AIC), ensuring the model accurately reflected the data’s underlying structure (Delignette-Muller & Dutang, 2015). Four independent Markov Chain Monte Carlo (MCMC) chains were run with 4,000 iterations each (2,000 warm-up); convergence was confirmed via R^ < 1.01 and sufficiently high effective sample sizes. Parameter estimates are reported as posterior medians with 95% credible intervals (CrI). Predictors were centered and scaled where appropriate to aid convergence.

## Results

### Extracellular K+ dynamics exhibit regional and circadian specificity

We first characterized how extracellular ([K^+^]ₒ) concentration changes in the hippocampus during and after standardized neuronal network activity, examining whether these dynamics differ between dorsal and ventral hippocampus across circadian time. We used a 10 Hz, 30-second stimulation of Schaffer collaterals to evoke robust [K^+^]ₒ release from activated neurons (Fig. 1A and B). This protocol was selected to ensure that the neuronal network response reached true peak amplitude and steady state before stimulus termination, enabling consistent comparison across experimental groups (see Methods).

**Figure 1.**
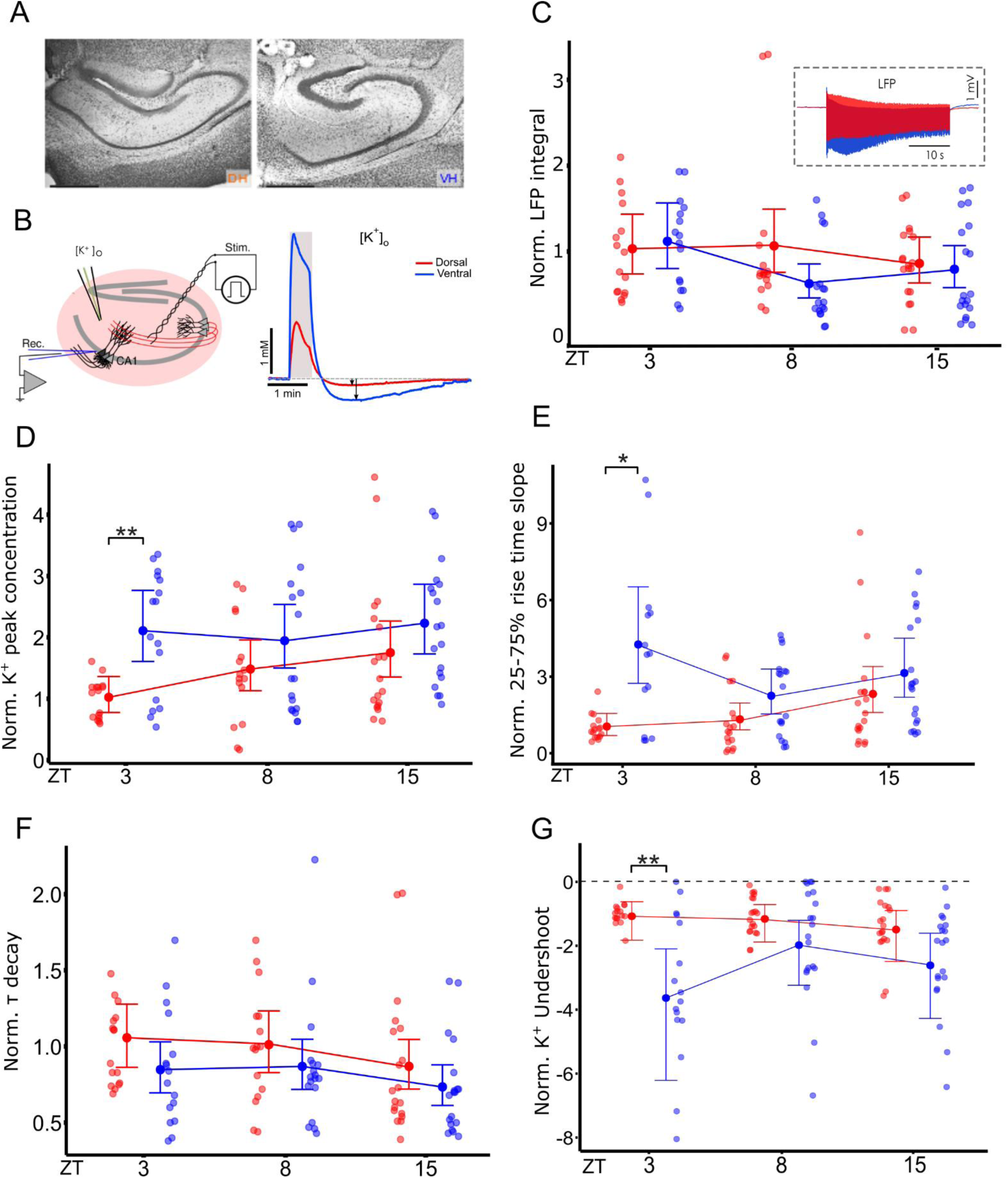
Network level extracellular K+ recording. **A)** Example of anatomical structures of dorsal (DH: red) and ventral (VH: blue) hippocampal slices. **B)** Schematic arrangement of stimulation of Schaffer Collateral (SC), recording of local field potential (LFP) from the *stratum oriens* (*s.o.*), and K+ recording [K^+^]ₒ from *stratum radiatum* (*s.r.*). Example trace of [K^+^]ₒ in DH (red) and VH (blue) in response to 10Hz30s stimulation. **C-G)** Normalized LFP integral, 25-75% rise time, peak amplitude, tau decay of monoexponentially fitted, and undershoot amplitude across DH and VH in three ZT3, ZT8, and ZT15 Analyses, based on a lognormal distribution model, are reported as posterior median estimates with 95% credible intervals. * *P* < 0.05, \*\**P* < 0.01. Statistical significance was calculated using two-sided unpaired Mann–Whitney U test. [DH_ZT3: n =16 (N = 8), DH_ZT8: n =16 (N= 8), DH_ZT5: n =19 (N = 10), VH_ZT3: n =16 (N=8), VH_ZT8: n =18 (N= 9), VH_ZT5: n =19 (N=10)]

### Synaptic strength is comparable across space and time

To ensure valid comparisons of [K^+^]ₒ transient properties, we first verified that evoked neuronal activity was equivalent between regions and time points. The magnitude of the local field potential (LFP) integral served as a quantitative proxy for network response intensity. Statistical analysis revealed no significant effects of hippocampal region (DH versus VH), Zeitgeber time (ZT), or their interaction on evoked LFP strength (Fig. 1C). This uniformity confirms the robustness of our stimulation paradigm and validates the assumption that any observed differences in [K^+^]ₒ dynamics reflect astrocyte properties rather than variations in neuronal input.

### Ventral hippocampus accumulates [K^+^]ₒ more rapidly and to greater amplitudes, particularly at early light phase

We assessed [K^+^]ₒ accumulation by analyzing the peak amplitude of [K^+^]ₒ transients. Permutation analysis of variance (ANOVA) shows a significant main effect of the regions only (*F*(1, 98) = 12.05, *permutation p* = .0010, ω^2^ = 0.0950), where the pool of VH samples shows 65% [95%CI 26%, 101%] larger [K^+^]ₒ peak amplitude than DH ones. Bayesian regression confirmed that this difference is predominantly driven by ZT3. At this time point, [K^+^]ₒ in the VH was 105% higher than in the DH [95% CrI: 40.5%, 200.4%]. In contrast, at ZT15, the DH exhibited a peak amplitude that was 71.6% greater than at ZT3 [95% CrI: 17.4%, 150.9%], suggesting that the transition from ZT3 (light cycle) to ZT15 (dark cycle) promotes greater [K^+^]ₒ accumulation in the DH (Fig. 1D).

Analysis of the slope of the 25–75% rise phase of [K^+^]ₒ transients revealed that VH reached peak [K^+^]ₒ 305.5% faster than the DH [95% CI: 124.8%, 646.3%] at ZT3. In the DH, the transition from ZT3 to ZT8 produced no clear change [27.1%, 95% CrI: −26.7%, 120.3%], whereas the transition to ZT15 resulted in a substantial increase in accumulation rate [122.6%, 95% CrI: 28.4%, 293.5%]. Importantly, the regional advantage of the VH diminished at later time points. Relative to its ZT3, the VH effect was attenuated by 41.7% at ZT8 [95% CrI: 19.0%, 93.1%] and by 33.3% at ZT15 [95% CrI: 14.9%, 72.7%], confirming a significant, negative interaction between region and time. (Fig. 1E).

Collectively, these results demonstrate that comparable neuronal network activity produces different extracellular [K^+^]ₒ dynamics between dorsal and ventral hippocampus, suggesting regional differences in astrocytic K^+^ buffering capacity. This regional divergence is most pronounced at the early light phase.

### [K^+^]ₒ clearance kinetics show similar rates despite regional differences in accumulation

To distinguish whether faster [K^+^]ₒ accumulation in VH reflects impaired [K^+^]ₒ removal or differential regulation at the rising phase of activity, we analyzed [K^+^]ₒ decay kinetics by fitting monoexponential decay (τ_decay_) to the falling phase of the transient. Permutation ANOVA revealed a significant main effect of region (F(1, 98) = 3.93, permutation p = 0.0496, ω² = 0.0278), with VH showing an overall 14.5% faster decay than DH [95% CI: -0.20%, 28.1%]. However, Bayesian regression analysis incorporating temporal variation did not identify statistically robust regional differences across time points. When analyzed within specific circadian phases, differences were modest and credible intervals crossed zero: at ZT3, DH exhibited 22.3% slower decay than VH [95% CrI: -5.9%, 50.2%]; at ZT8 and ZT15, differences were 15.2% [95% CrI: -14%, 41.5%] and 17.0% [95% CrI: -11%, 42.1%], respectively (Fig. 1F).

This paradoxical finding – accelerated [K^+^]ₒ accumulation in VH despite similar or faster clearance rates – indicates that differential [K^+^]ₒ dynamics are driven by regional difference at the rising phase of the transient rather than the decay phase, suggesting that distinct buffering mechanisms may be preferentially engaged during accumulation versus recovery.

### Ventral hippocampus displays enhanced putative Na^+^/K^+^-ATPase activity specifically at early light phase

The similar decay kinetics between regions despite higher [K^+^]ₒ accumulation in VH prompted us to examine the recovery phase. After prolonged stimulation (10 Hz, 30 s), [K^+^]ₒ return to baseline is followed by an undershoot (Fig. 1B, right panel, black arrows), a transient decrease of K^+^ concentration below baseline level that reflects Na^+^/K^+^-ATPase activity (Chever et al., 2010; D’Ambrosio et al., 2002). Analysis of undershoot amplitude revealed significant regional effects and complex temporal dynamics: permutation ANOVA revealed a significant effect of region (F(1, 105) = 26.65, permutation p = 0.0001, ω² = 0.180) as well as significant interaction of time with region (F(2, 105) = 3.074, permutation p = 0.0482, ω² = 0.0290). Bayesian regression analysis revealed that at ZT3, VH showed a 239% increase in undershoot amplitude compared to DH [95% CrI: 54%, 624%]. While VH maintained elevated undershoot amplitudes at ZT8 and ZT15, the magnitude of this regional difference was attenuated at these later timepoints [ZT8: -49%, 95% CrI: -82%, +42%; ZT15: -48%, 95% CrI: -81%, +48%] (Fig. 1G).

These findings suggest that VH astrocytes display enhanced Na^+^/K^+^-ATPase activity, particularly pronounced at the early light phase. Despite this elevated pump activity, overall [K^+^]ₒ clearance rates remain unchanged, suggesting that the enhanced pump activity primarily reflects the greater [K^+^]ₒ accumulation in VH (which provides greater driving force for pump activation) rather than fundamentally altering the rate constant of [K^+^]ₒ removal. This raises the question: what cellular mechanisms limit [K^+^]ₒ buffering capacity in the ventral hippocampus?

### Cellular basis of regional [K^+^]ₒ buffering differences: a role for reduced Kir4.1 function in ventral astrocytes

#### Astrocytic Gap Junction Coupling Shows Regional and Temporal Variation

Potassium redistribution through astrocytic networks depends on gap junction-mediated coupling, which expands the effective volume available for [K^+^]ₒ uptake. We examined whether astrocytic coupling—measured by quantifying the number of biocytin-labeled cells connected to a patched astrocyte—exhibits regional and circadian variation (Fig. 2A). Across all time points, VH slices showed a trend toward more coupled astrocytes than DH [mean difference: 13 cells, 95% CI: -4, 30], but this difference did not reach statistical significance in permutation ANOVA (F(2, 63) = 2.75, permutation p = 0.0686, ω² = 0.0483). However, Bayesian regression analysis revealed significant regional and temporal effects. Specifically, at ZT3, VH slices exhibited 39 more coupled cells than DH [95% CrI: +15, +69], corresponding to a 36.3% increase [95% CrI: 8.3%, 70.0%]. This regional difference was absent at ZT8. At ZT15, DH coupling increased by 27% compared to its ZT3 value [95% CrI: 1%, 60%], while VH coupling declined by 16.2% [95% CI: 6.8%, 52.3%] (Fig. 2A and B).

**Figure 2.**
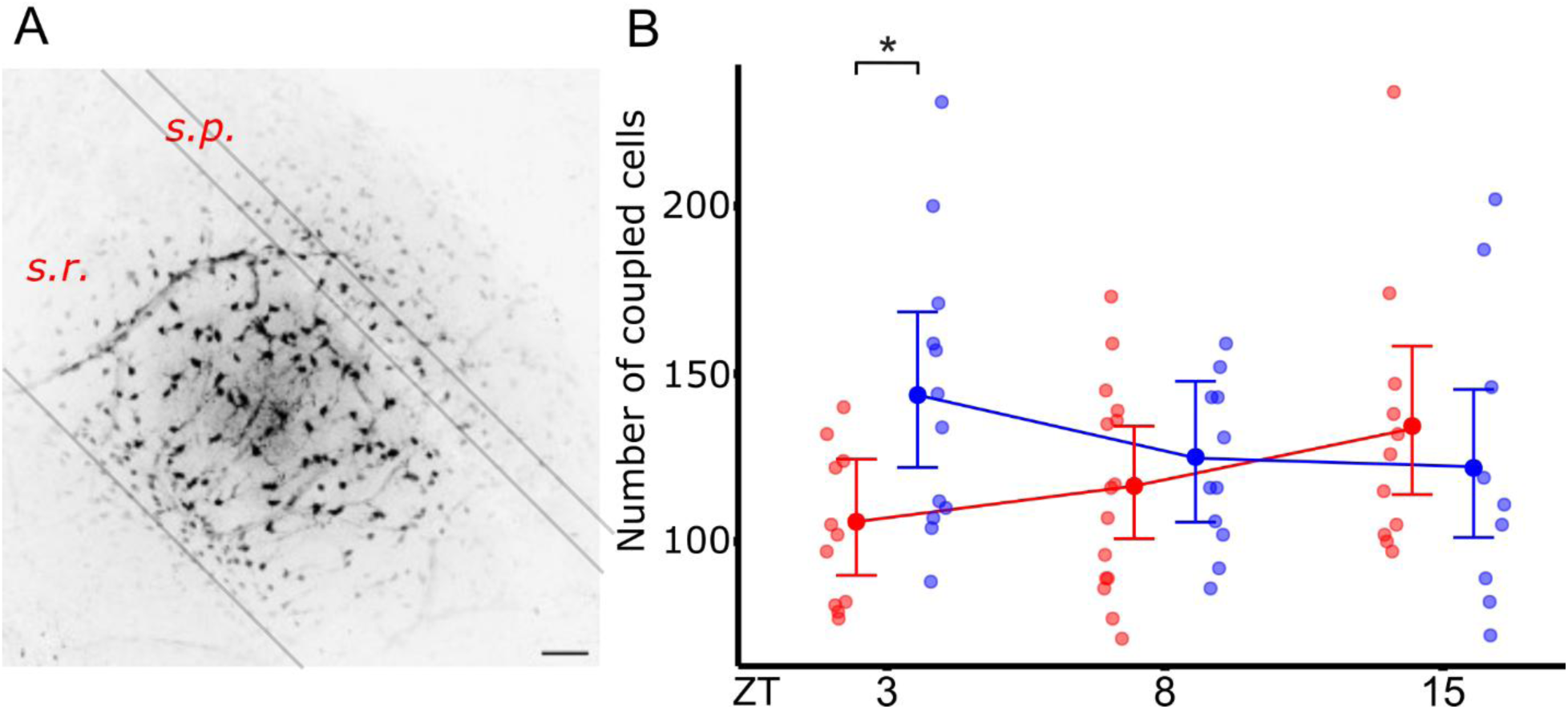
Gap junction coupling across hippocampal regions and time of day. **A)** Inverted example image of the coupled astrocytes in a hippocampal slice, as obtained after biocytin diffusion from a single cell, held in whole-cell voltage clamp for 20 mins. Scale bar is equal to 50 µm. **B)** Summary of the number of biocytin-coupled cell across DH and VH in different time points (ZT3, 8, 15). Analyses, based on a lognormal distribution model, are reported as posterior median estimates with 95% credible intervals. *p < 0.05 Mann–Whitney U test; *s.p., Stratum pyramidale; s.r., stratum radiatum*. [DH_ZT3: n =11 (N = 6), DH_ZT8: n =15 (N= 6), DH_ZT5: n =11 (N = 6), VH_ZT3: n =12 (N=7), VH_ZT8: n =11 (N= 7), VH_ZT5: n =9 (N=6)]

Enhanced gap junction coupling in VH at ZT3 would be expected to facilitate [K^+^]ₒ redistribution and thus improve buffering capacity. The fact that VH nevertheless shows greater [K^+^]ₒ accumulation at this timepoint suggests that reduced K^+^ conductance mechanisms (rather than impaired redistribution) are the primary determinant of regional differences.

#### Ventral hippocampal astrocytes display reduced Kir4.1-mediated conductance across all circadian phases

Kir4.1 channels mediate the primary K^+^-dependent astrocytic conductance and are central to spatial buffering, particularly during the rising phase of [K^+^]ₒ transients when astrocytic depolarization drives K^+^ influx through these channels (Larsen et al., 2014; Seifert et al., 2009). To directly assess Kir4.1 function, we performed whole-cell patch-clamp recordings (Fig. 3A) in the presence of gap junction blocker (MFA) and neuronal transmission blockers (TTX, CNQX, D-AP5), isolating Kir4.1-specific currents via subsequent Ba²^+^ application (Wallraff et al., 2006). Indeed, application of MFA was demonstrated to be effective in uncoupling astrocytes, as shown by the number of dye-coupled cells (Fig. 3B-C).

**Figure 3.**
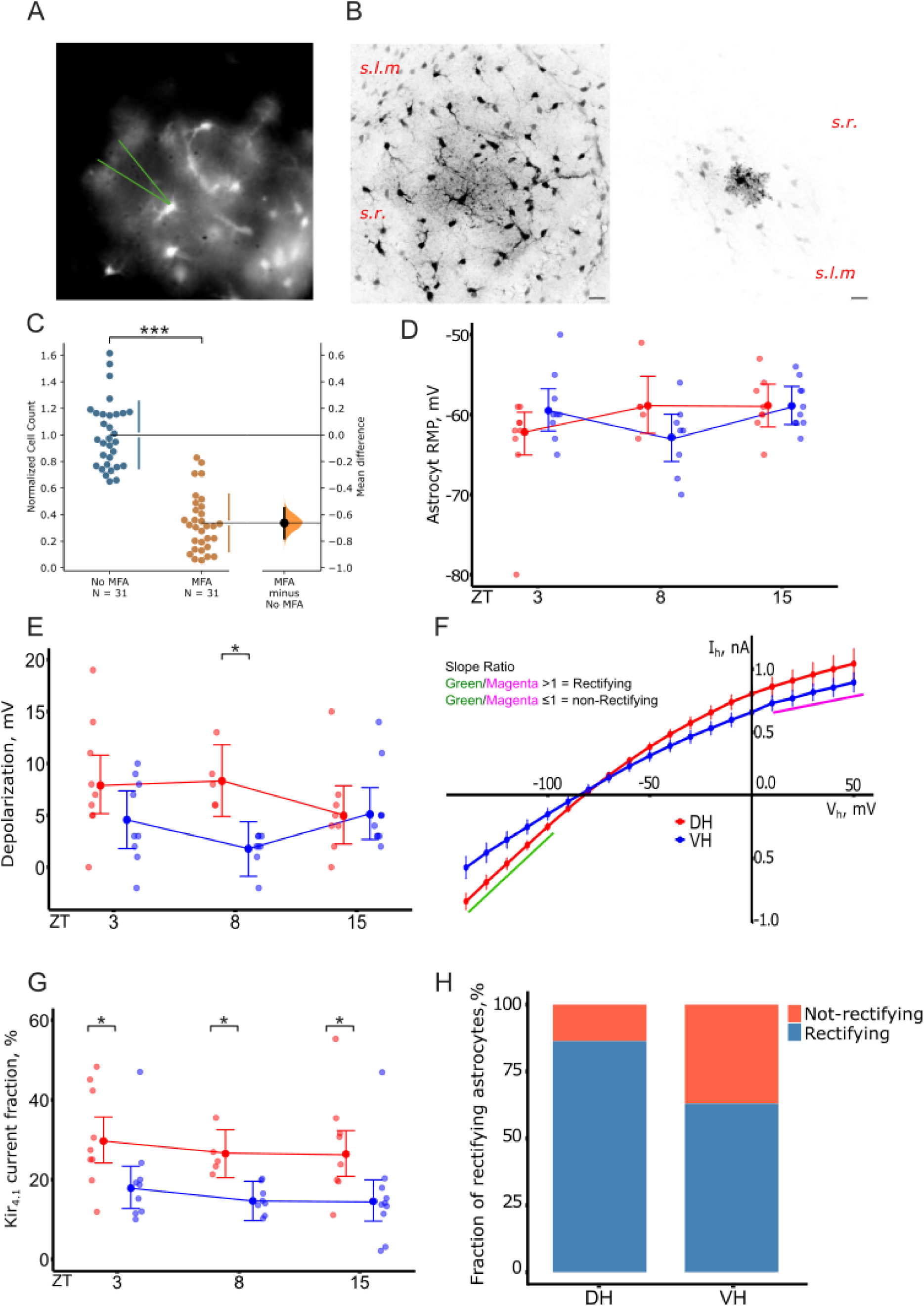
Regional differences across regions and time. **A)** Wide-field image of stained astrocytes with SR101. **B)** Left: example image of biocytin stained astrocytes in absence of MFA versus incubated in MFA (right). *S.r., stratum radiatum; s.l.m., stratum lacunosum moleculare*. Scale bar is equal to 20 µm. **C)** statistical summary of the coupled cell number in both groups. Addition of MFA results in an average reduction of -86 [95%CI -95, -72], p < .00001. **D)** Initial resting membrane potential (RMP; in mV) of the patched astrocytes across DH and VH in ZT3, ZT8, and ZT15. **E)** Ba2+-induced depolarization across the regions and time. No significant effect of region and time was observed, except in an isolated incident of ZT8. **F)** Ba2+-sensitive current obtained by subtracting membrane currents elicited in whole-cell configuration (from -140 mV to +50 mV, 10 mV increment; holding potential = −70 mV) from the baseline: DH (red) and VH (blue). The rectification of measurements are based on the barium-sensitive currents. The IV curve slope of -140 to -100mV interval (green) was divided to the slope of 10mV to 50mV (magenta), and rectification profile of the tested cell was determined given the ratios. **G**) ) Kir4.1 current across the regions and time of day. Significant main effect of region was observed without the influence of time. Analyses are reported as posterior median estimates with 95% credible intervals. **H)** Proportion of rectifying astrocytes (blue) versus non-rectifying (orange) across DH and VH. [DH_ZT3: n = 9 (N = 7), DH_ZT8: n = 5 (N= 3), DH_ZT5: n =8 (N = 6), VH_ZT3: n =9 (N=6), VH_ZT8: n =8 (N= 5), VH_ZT5: n =10 (N=5)] * *P* < 0.05, \*\*\**P* < 0.001. Statistical significance was calculated using two-sided unpaired Mann–Whitney U test

Resting membrane potential did not vary significantly between regions or across time (Fig. 3D). Application of Ba²^+^ caused an average depolarization of 5.5 ± 0.64 mV consistent with blockade of inward-rectifying currents, confirming effective channel blockade. Permutation ANOVA revealed a significant main effect of region on Ba²^+^-induced depolarization (F(1, 43) = 8.26, permutation p = 0.0051, ω² = 0.124), with DH astrocytes depolarizing 3.2 mV more than VH astrocytes following Ba²^+^ application [95% CI: 0.90, 5.70], indicating higher functional Kir4.1 expression in DH (Fig. 3E).

To more directly quantify Kir4.1-mediated conductance, we analyzed Ba²^+^-sensitive current at -130 mV holding potential, a voltage optimal for Kir4.1 channel activity (Seifert et al., 2009). Permutation ANOVA revealed a significant effect of region (F(1, 43) = 13.30, permutation p = 0.0006, ω² = 0.209). Bayesian regression confirmed a significant main effect of region, with no effect of time or region-time interaction. On average, Ba²^+^-sensitive conductance represents 29% of total transmembrane current in DH astrocytes compared to 17% in VH, with a mean difference of 12% [95% CI: 5.8%, 17.6%] (Fig. 3F-G). This reduction in functional Kir4.1 conductance in VH relative to DH persists across all examined circadian phases, suggesting that regional differences in Kir4.1 function are stable properties of these astrocytes rather than acutely regulated.

This reduced Kir4.1-mediated conductance in VH astrocytes provides a mechanistic explanation for the observed rapid and pronounced K^+^ accumulation in this region: with lower K^+^ conductance, astrocytes can passively take up less K^+^ in response to depolarization, resulting in higher [K^+^]ₒ despite the presence of enhanced gap junction coupling that could support redistribution.

We noticed that in some astrocytes the IV relationship of Ba^2+^-sensitive current was linear without rectification. We checked whether such astrocytes are more frequent in one of the explored regions. Current rectification was assessed by dividing I-V curve slopes between – 140 to –90 mV by that of 0 to +50 mV voltage ranges. Cells with slope ratios >1 were classified as rectifying. Fisher’s exact test revealed no statistically significant association between region and rectification status (p = 0.104), though the estimated odds ratio of 0.28 [95% CI: 0.04, 1.31] suggested a non-significant trend toward greater rectification in DH astrocytes (Fig. 3H). Our findings indicate that Kir4.1-mediated currents are regionally regulated, with higher functional expression in the DH compared to VH but are not modulated by circadian time under these experimental conditions. This difference partly explains why, at the same rate of potassium release, its accumulation is faster and greater in VH than in DH.

### Kir4.1 protein expression exhibits circadian dynamics distinct between dorsal and ventral hippocampus

To investigate whether regional differences in Kir4.1 current reflect corresponding differences in protein abundance, and to assess circadian regulation of Kir4.1 expression, we performed double immunolabeling for GFAP (astrocyte marker) and Kir4.1 on hippocampal sections across the regions (Fig. 4A-B) as well as time points.

**Figure 4.**
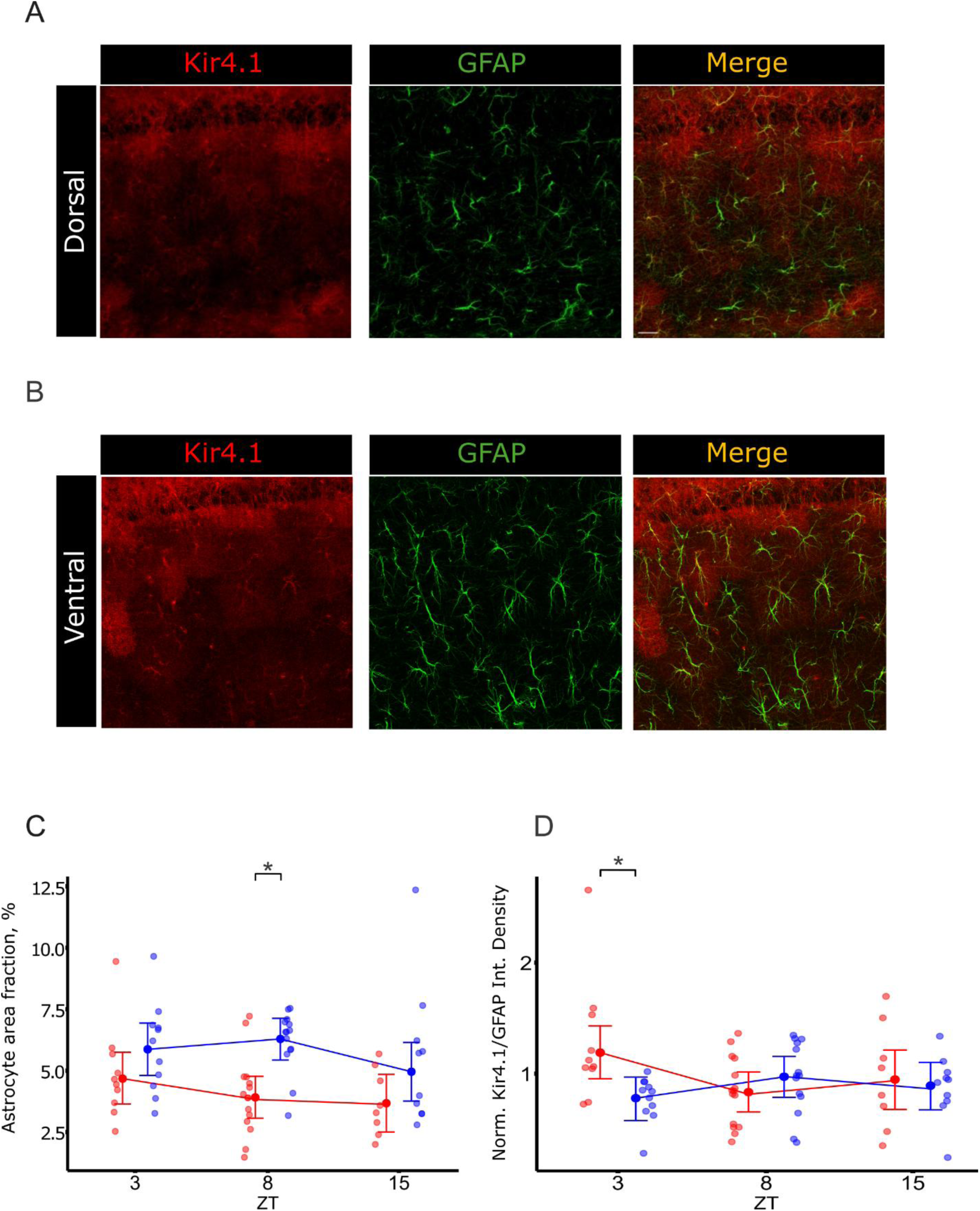
Regional differences in Kir4.1 expression along the dorsoventral axis depends on time. **A)** Representative examples of confocal images of immunofluorescent labeling of Kir4.1 (red) and astrocyte marker glial fibrillary acidic protein (GFAP, green), and DAPI (blue) for DH (Left) and VH (Right), at 20x magnification (scale bar = 100 µm). **B)** Higher magnification of confocal images of Kir4.1 (red) and astrocyte marker glial fibrillary acidic protein (GFAP, green), and the merging of the two. Images were taken from the s.r. **C)** Fraction of GFAP positive area with Kir4.1 protein is significantly smaller in DH -1.72% [95%CI -2.6%, -0.88%], and this effect is driven by ZT8 -2.21% [95%CI -3.12%, -1.11%]. **D)** Quantification of Kir4.1 integrated density normalized to GFAP. The relationship was only significant in ZT3, where VH showed -46% (less) than DH [95%CI -74%, -19%]. Analyses, based on a student t-distribution model, are reported as posterior median estimates with 95% credible intervals. * *P* < 0.05. Statistical significance was calculated using two-sided unpaired Mann–Whitney U test [DH_ZT3: n = 11 (N = 1), DH_ZT8: n = 16 (N= 1), DH_ZT5: n =9 (N = 1), VH_ZT3: n =12 (N=1), VH_ZT8: n =15 (N= 1), VH_ZT5: n =11 (N=1)]

Analysis of GFAP-positive area fraction revealed a main effect of region (F(1, 62) = 13.48, permutation p = 0.0009, ω² = 0.157), although Bayesian analysis detected no statistically robust effects (Fig. 4C). To account for regional differences in astrocytic Kir4.1 expression, Kir4.1 integrated density was normalized to GFAP-positive area (astrocyte-specific quantification). Values were further normalized within each immunolabeling batch to DH samples to control batch variability.

Permutation ANOVA revealed a significant region × time interaction (F(2, 62) = 4.14, permutation p = 0.0119, ω² = 0.092). Bayesian regression analysis revealed nuanced circadian regulation: at ZT3 (early light), DH showed significantly higher Kir4.1 levels than VH [39.3% increase, 95% CrI: 14.8%, 56.8%]. The transition from ZT3 to ZT8 (late light) was marked by significant decline in DH Kir4.1 expression [34.3% decrease, 95% CrI: 10.4%, 51.8%], whereas VH showed the opposite pattern with a 24.4% increase [95% CrI: 22.1%, 197.4%], resulting in substantial convergence of expression levels. No reliable regional differences were detected at ZT15 (early dark) (Fig. 4D).

Importantly, these circadian changes in Kir4.1 protein abundance do not translate into circadian modulation of Kir4.1-mediated currents (which showed only regional, not temporal, variation in functional assays). This dissociation between protein expression dynamics and current magnitude suggests complex post-translational regulation of Kir4.1 channel function.

#### Integration and mechanistic synthesis

Our results reveal a multiscale organizational principle for astrocytic K^+^ buffering in hippocampus: regional and circadian factors converge to determine buffering capacity through complementary mechanisms operating at different phases of [K^+^]ₒ transients (Fig. 5).

**Figure 5.**
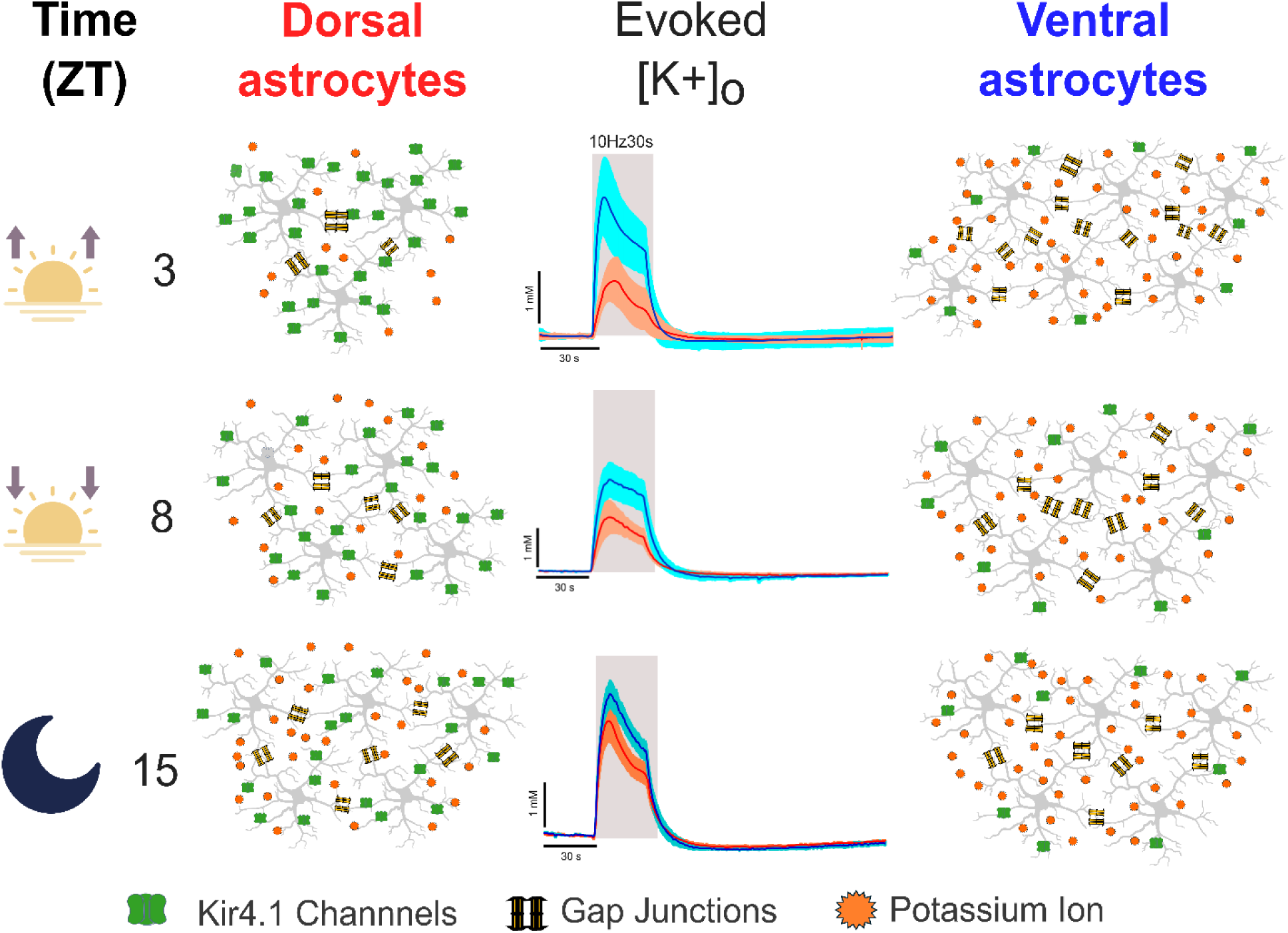
Graphical summary of the results.

At the early light phase (ZT3), the ventral hippocampus exhibits a state of heightened extracellular K^+^ accumulation characterized by: 1) reduced Kir4.1-mediated K^+^ conductance (41% lower than DH) that persists across all times, 2) enhanced gap junction coupling (35% more coupled astrocytes) that could support redistribution, 3) elevated Na^+^/K^+^-ATPase-dependent undershoot (239% greater than DH) indicating compensatory pump activation. These properties produce rapid and pronounced [K^+^]ₒ accumulation despite the presence of potentially advantageous gap junction coupling. The putative enhanced Na^+^/K^+^-ATPase activity likely reflects activation by the greater local [K^+^]ₒ accumulation and astrocytic depolarization, coupled with potentially higher pump expression. Thus, at ZT3, ventral astrocytes are limited in their capacity to prevent [K^+^]ₒ accumulation during the rising phase.

As the circadian cycle progresses through the light phase (ZT3 to ZT8 to ZT15): 1) Kir4.1 protein expression increases in VH and decreases in DH, partially equalizing regional differences by ZT15, 2) gap junction coupling normalizes between regions, 3) Na^+^/K^+^-ATPase compensatory activity diminishes.

By the early dark phase (ZT15), the circadian pattern reverses, with DH [K^+^]ₒ accumulation increasing relative to ZT3, suggesting that dorsal hippocampal astrocytes are subject to time-dependent suppression of buffering capacity during the light phase, with restoration during the dark phase. The nature of this circadian suppression remains to be determined but may reflect circadian regulation of Kir4.1 trafficking or channel modulation by clock-controlled signaling pathways.

## Discussion

This study addresses the question of spatial and temporal heterogeneity in astrocytic function. Using integrated functional, electrophysiological, and molecular approaches, we demonstrate that astrocytic K^+^ buffering in the hippocampus is jointly regulated by regional identity and circadian phase, with important implications for understanding network excitability, seizure susceptibility, and the temporal organization of brain states.

### Astrocytes show regional specialization but do not simply mirror neuronal excitability patterns

Our findings reveal that ventral hippocampal astrocytes possess fundamentally different [K^+^]ₒ buffering properties compared to their dorsal counterparts. At the early light phase (ZT3), when regional differences are most pronounced, VH accumulates [K^+^]ₒ more rapidly and to higher peak amplitudes than DH despite equivalent neuronal network activity. This regional divergence reflects intrinsic differences in astrocyte physiology rather than variations in neuronal input, establishing that astrocytic heterogeneity is a primary organizational principle of hippocampal circuits.

However, astrocytic regional specialization does not simply track neuronal excitability in a compensatory manner. The VH is known to exhibit higher intrinsic neuronal excitability and greater seizure susceptibility than the DH (Debski et al., 2020; Dougherty et al., 2012; Papatheodoropoulos et al., 2005). If astrocytes were optimized to compensate for regional differences in neuronal activity, one would predict enhanced [K^+^]ₒ buffering capacity in VH to counterbalance its elevated excitability. Instead, we observe the opposite: VH astrocytes display reduced Kir4.1-mediated conductance (12% lower than DH), resulting in greater [K^+^]ₒ accumulation during neuronal activity. This counterintuitive arrangement suggests that astrocytic properties are not configured to suppress regional differences in excitability but rather may amplify or enable them, supporting the functionally distinct roles mediated by dorsal and ventral hippocampus in cognition and behavior.

### The paradox of enhanced accumulation with preserved clearance: differential mechanisms at rising vs. falling phases

A central finding of this study is the paradoxical observation that VH exhibits both faster [K^+^]ₒ accumulation and similar (or slightly faster) clearance rates compared to DH. This apparent contradiction is resolved by recognizing that distinct buffering mechanisms dominate during different phases of the K^+^ transient, with Kir4.1 channels playing a central role during accumulation and Na^+^/K^+^-ATPase predominating during recovery.

In this study, Na^+^/K^+^ ATPase activity was quantified using the amplitude of the post-stimulus K^+^ undershoot rather than by pharmacological pump inhibition because it does not selectively reduce [K^+^]ₒ clearance. Indeed, the pump inhibition depolarizes neurons, alters firing patterns, and modifies synaptic transmission (T. R. Anderson et al., 2010), thereby changing K+ release during Schaffer collateral stimulation. Consequently, [K^+^]ₒ transients recorded after pump blockade reflect a mixture of impaired clearance and altered K^+^ release, which cannot be disentangled. In contrast, the [K^+^]ₒ undershoot that follows a defined activation epoch arises from activity-dependent stimulation of the Na^+^/K^+^ ATPase and reflects the excess pump-mediated uptake of K^+^ once neuronal firing has ceased. For these reasons, undershoot analysis offers a more specific and interpretable functional measure of Na^+^/K^+^ ATPase activity in our experimental condition.

During the rising phase of neuronal activity, when K^+^ is actively released from depolarizing neurons, [K^+^]ₒ elevation depolarizes nearby astrocytes, driving passive K^+^ influx through Kir4.1 channels down the electrochemical gradient (Larsen & MacAulay, 2014). This Kir4.1-mediated uptake represents the fastest way to limit [K^+^]ₒ accumulation. Our electrophysiological data demonstrate that VH astrocytes possess significantly reduced Ba²^+^-sensitive (Kir4.1-mediated) conductance compared to DH astrocytes, a decrease that persists across all circadian phases examined. This reduced conductance directly explains why VH accumulates [K^+^]ₒ more rapidly: with fewer functional Kir4.1 channels, astrocytes cannot buffer K^+^ as efficiently during the rising phase, resulting in higher extracellular concentrations.

In contrast, during the decay phase after neuronal activity ceases, [K^+^]ₒ clearance is dominated by active Na^+^/K^+^-ATPase pumping rather than passive Kir4.1-mediated influx (D’Ambrosio et al., 2002; MacAulay, 2020). Our observation that VH exhibits a dramatically enhanced [K^+^]ₒ undershoot (239% larger than DH at ZT3)—a signature of robust Na^+^/K^+^-ATPase activity—indicates that pump-mediated clearance is elevated in VH. Importantly, this enhanced pump activity likely reflects a compensatory response to the greater [K^+^]ₒ accumulation itself: higher [K^+^]ₒ and greater astrocytic depolarization both stimulate Na^+^/K^+^-ATPase activity (Wang et al., 2012). Thus, the similar decay kinetics between regions emerge not from equivalent intrinsic clearance capacity but from activity-dependent upregulation of pumping in VH that compensates for its reduced Kir4.1 function.

This mechanistic model—reduced Kir4.1-mediated uptake during accumulation, compensated by enhanced pump-mediated clearance during recovery—reconciles the paradoxical K^+^ dynamics and highlights the phase-specific contributions of different buffering mechanisms.

### Gap junction coupling: enhanced connectivity cannot overcome reduced conductance

An unexpected finding was that VH astrocytes exhibit enhanced gap junction coupling at ZT3 (35% more coupled cells than DH), precisely when and where K^+^ accumulation is greatest. At first glance, this enhanced coupling appears maladaptive: expanded astrocytic networks should increase the effective volume available for K^+^ redistribution and thus improve buffering capacity (Wallraff et al., 2006). Yet VH accumulates more [K^+^]ₒ.

This paradox highlights an important distinction between network connectivity (assessed by gap junction coupling) and cellular K^+^ conductance (determined primarily by Kir4.1 expression). Enhanced coupling expands the astrocytic syncytium, but K^+^ must first enter astrocytes through Kir4.1 channels before it can be redistributed through the network. With Kir4.1 conductance reduced by 12% in VH astrocytes, the membrane represents a bottleneck that limits K^+^ uptake regardless of how extensive the downstream network may be. Thus, gap junction coupling appears to represent a partial compensatory adaptation—expanding the potential redistribution capacity—but this adaptation cannot fully overcome the primary deficit in membrane conductance.

This interpretation suggests that Kir4.1 channel function, rather than gap junction coupling, is the rate-limiting step for K^+^ buffering during active neuronal stimulation in VH.

### Circadian regulation superimposed on stable regional identity

Our temporal analysis reveals that circadian phase modulates—but does not abolish—the stable regional differences in K^+^ buffering. Two key parameters exhibit no circadian variation: (1) the time constant of K^+^ decay, which remains similar between regions and constant across time, and (2) Ba²^+^-sensitive Kir4.1 current, which shows consistent regional differences (lower in VH) at all timepoints despite circadian changes in Kir4.1 protein expression.

The stability of Kir4.1 current despite dynamic protein expression points to complex post-translational regulation. At ZT3, DH exhibits 41% higher Kir4.1 protein than VH; by ZT8, this regional difference disappears due to reduced expression in DH and increased expression in VH. Yet functional Kir4.1 currents remain regionally differentiated throughout the day. This dissociation suggests that Kir4.1 channel activity is regulated not only by protein abundance but also by trafficking to the membrane, phosphorylation state, interaction with scaffolding proteins, or lipid microenvironment—mechanisms that may themselves be under circadian control and differ between regions.

The most pronounced circadian effect is observed in the convergence of multiple parameters at ZT3 versus later timepoints. At early light phase, the regional divergence between DH and VH is maximal: K^+^ peak amplitude differs 2.1-fold, gap junction coupling differs by 35%, and Kir4.1 protein expression differs by 41%. By ZT8 and ZT15, these differences attenuate or reverse. This temporal pattern suggests that early light phase—corresponding to the rest period in nocturnal rodents—may be a time when regional specialization is most important, possibly reflecting distinct roles for dorsal and ventral hippocampus in sleep-dependent memory consolidation, sharp-wave ripple generation, or metabolic recovery (Pronier et al., 2023).

### Functional implications: regional K^+^ dynamics shape circuit computations

The distinct K^+^ buffering properties of DH and VH astrocytes have important implications for how these regions process information and contribute to hippocampal function. Extracellular K^+^ accumulation exerts multiple effects on neuronal excitability, synaptic transmission, and network oscillations, making astrocytic control of [K^+^]ₒ a powerful regulator of circuit dynamics.

Ventral hippocampus: permissive K^+^ accumulation enables synchronized network events but imposes low-pass filtering. The rapid and pronounced [K^+^]ₒ accumulation in VH has both facilitatory and inhibitory consequences. Elevated [K^+^]ₒ depolarizes neurons toward action potential threshold, enhancing NMDA receptor activation by relieving Mg²^+^ block and increasing the spatial range of glutamate spillover (Shih et al., 2013). These effects promote network synchronization and may facilitate the generation of hippocampal sharp-wave ripples (SWRs)—brief, high-frequency oscillations essential for memory consolidation. Indeed, VH slices more readily generate spontaneous SWRs than DH slices (Schlingloff et al., 2014), and CA3 pyramidal neurons in VH possess stronger recurrent connectivity and higher intrinsic excitability (Sun et al., 2020), properties that synergize with permissive K^+^ accumulation to support synchronized discharges.

However, sustained K^+^ elevation during prolonged activity also inactivates voltage-gated Na^+^ channels and reduces the driving force for K^+^ efflux through voltage-gated K^+^ channels,

leading to synaptic depression and reduced responsiveness to repetitive stimulation (Meeks & Mennerick, 2004). This creates a functional low-pass filter, where VH responds robustly to initial or low-frequency inputs but saturates during sustained high-frequency activity. Such filtering may prevent runaway excitation and protect against seizure propagation, while simultaneously biasing VH toward processing sparse, salient events rather than sustained information streams.

Dorsal hippocampus: tight K^+^ control preserves high-fidelity transmission and supports frequency-dependent facilitation. In contrast to VH, the strong Kir4.1-mediated buffering in DH maintains lower [K^+^]ₒ during neuronal activity, preserving neuronal excitability and enabling synaptic facilitation at frequencies between 10-50 Hz—precisely the range that encodes behaviorally relevant information (Papaleonidopoulos et al., 2017). Lower K^+^ accumulation prevents synaptic depression, supporting sustained information transfer and communication with neocortical targets (Moreno et al., 2016). This configuration suits DH for high-pass filtering: reliable transmission of sustained, structured activity patterns required for spatial navigation, episodic memory encoding, and cognitive map formation (Fanselow & Dong, 2010).

The metabolic consequences of these regional differences are consistent with recent findings that VH relies more heavily on aerobic glycolysis—generating cytosolic ATP rapidly but inefficiently—compared to the oxidative phosphorylation favored in DH (Brancati et al., 2021). The elevated Na^+^/K^+^-ATPase activity required to clear excess K^+^ in VH imposes substantial energetic demands that may be preferentially met by glycolytic metabolism.

### Pathological implications: regional K^+^ dysregulation and epilepsy

The reduced K^+^ buffering capacity of VH astrocytes provides a mechanistic framework for understanding the well-established vulnerability of VH to seizures. Evidence from human temporal lobe epilepsy (TLE) and animal models identifies VH as the primary seizure initiation zone (Babb et al., 1984; Bernasconi et al., 2003; Buckmaster et al., 2022; Toyoda et al., 2013). Our findings suggest that intrinsic differences in astrocytic K^+^ buffering—present even in healthy tissue—predispose VH to hyperexcitability.

During pathological conditions such as epilepsy, when neuronal activity is abnormally intense and synchronized, the limited Kir4.1 capacity in VH would result in excessive [K^+^]ₒ accumulation, reaching levels (>10 mM) known to be associated with or even trigger seizures (de Curtis et al., 2018; Enger et al., 2015). Furthermore, epileptic tissue exhibits downregulation of Kir4.1 expression and function (Bataveljic et al., 2024; Romanos et al., 2020), exacerbating the baseline deficit in VH and creating a positive feedback loop, i.e. reduced buffering results in greater K^+^ accumulation, which enhances excitability and thus more intense neuronal firing, favoring further K^+^ release.

The circadian modulation of K^+^ buffering we observe also provides a potential mechanism for the circadian rhythmicity of seizure occurrence documented in epilepsy patients and animal models (B.-L. Chang et al., 2018; Proix et al., 2021; Quigg et al., 2000). Seizure susceptibility peaks during specific circadian phases, and our data suggest that times of maximal regional divergence in K^+^ buffering (e.g., ZT3) may correspond to windows of heightened vulnerability in VH. Conversely, circadian phases when regional differences attenuate may represent periods of reduced seizure risk. However, given the extensive reorganization of circuits in epilepsy, it will be important to assess all the above-described properties in experimental epilepsy.

### Dissociation between Kir4.1 protein expression and functional current: post-translational complexity

One of the most intriguing findings is the dissociation between Kir4.1 protein expression dynamics (which vary significantly across circadian time) and functional Kir4.1-mediated currents (which show stable regional differences but no temporal variation). This dissociation implies that channel function is regulated at multiple levels beyond simple protein abundance.

Several post-translational mechanisms could account for this complexity. Kir4.1 channels must traffic to the plasma membrane to be functional, and this trafficking is regulated by interactions with scaffolding proteins such as dystrophin-associated protein complex and α-syntrophin (reviewed in (Olsen & Sontheimer, 2008)). Circadian changes in trafficking machinery or cytoskeletal organization could alter the proportion of synthesized channels that reach the membrane. Additionally, Kir4.1 activity is modulated by phosphorylation, pH, and membrane lipid composition (Hibino et al., 2010)—all of which exhibit circadian oscillations (McCauley et al., 2020; Ryu et al., 2024). Finally, Kir4.1 channels interact with other membrane proteins in macromolecular complexes, and changes in these interactions could alter channel open probability or conductance without changing total protein levels.

The persistence of regional differences in functional current despite convergence of protein expression also suggests that DH and VH astrocytes may differ in their complement of regulatory proteins or post-translational modifications. Identifying these region-specific regulatory mechanisms represents an important direction for future research and could reveal novel targets for modulating astrocytic K^+^ buffering in disease states.

### Comparative context: astrocytic heterogeneity as a general principle

Our findings contribute to a growing recognition that astrocytic heterogeneity is a fundamental organizational principle across the brain. Astrocytes differ between brain regions in molecular marker expression, morphology, calcium signaling dynamics, glutamate uptake capacity, and contribution to synaptic function (reviewed in (Khakh & Deneen, 2019)). Within the hippocampus, dorsoventral differences in astrocytic gene expression, morphology, and calcium dynamics have been documented (Dong et al., 2009; Jinno, 2011; Ryu et al., 2024; Thompson et al., 2008), but the functional consequences of this heterogeneity for circuit physiology have remained unclear.

Our demonstration that astrocytic K^+^ buffering—a core homeostatic function—exhibits pronounced regional and temporal variation establishes that astrocytic heterogeneity is not merely a descriptive phenomenon but has direct, measurable impacts on network excitability and information processing. This principle likely extends beyond the hippocampus to other brain structures where astrocytic properties may be tuned to local circuit demands.

Importantly, our findings challenge the traditional view of astrocytes as passive, uniform support cells that maintain brain homeostasis in a spatially and temporally invariant manner. Instead, astrocytes emerge as active, heterogeneous participants in circuit function whose properties are dynamically regulated in space and time. This has implications for understanding both normal brain function and the pathophysiology of neurological disorders.

### Reconciling slice physiology with in vivo function: caveats and considerations

Several important caveats limit the interpretation of our findings. First, our measurements were performed in acute brain slices, which lack the intact vascular system, interstitial fluid flow, and long-range connectivity present in vivo. K^+^ clearance mechanisms that rely on vascular uptake or long-distance redistribution through white matter tracts may be compromised in slices, potentially altering the relative contributions of different buffering mechanisms. Evidence suggests that gap junction-mediated spatial buffering contributes more to K^+^ clearance in vivo than in slices, raising the possibility that the enhanced gap junction coupling we observe in VH may be more functionally significant in intact brain than our slice experiments suggest (Breithausen et al., 2020; Cooper et al., 2025).

Second, patch-clamp approach in astrocytes samples a small radius of somata (Zhou et al., 2021), which may not fully represent the properties of fine perisynaptic processes where K^+^ buffering most directly impacts synaptic function. Recent advances in voltage imaging reveal that distal astrocytic processes experience larger depolarizations in response to neuronal activity than somata do (Armbruster et al., 2022), and spatial compartmentalization of ion channels may create local K^+^ buffering microdomains that our somatic recordings cannot detect. Additionally, we used GFAP immunolabeling to define astrocytic regions for Kir4.1 quantification, but GFAP labels only ∼15% of astrocyte volume (Bushong et al., 2002), biasing our analysis toward the most GFAP-rich cellular compartments and underestimating Kir4.1 in fine processes.

Third, our sampling of three circadian timepoints (ZT3, ZT8, ZT15) provides valuable initial insight into temporal regulation but may miss peak effects occurring at other phases. Higher temporal resolution studies examining more timepoints throughout the 24-hour cycle would provide a more complete picture of circadian dynamics.

Despite these limitations, our findings reveal fundamental principles of astrocytic K^+^ regulation that likely persist—and may be amplified—in vivo. The regional differences we document are substantial (2-fold differences in K^+^ peak amplitude, 41% difference in Kir4.1 current) and consistent across multiple experimental approaches, suggesting they reflect robust biological properties rather than experimental artifacts.

### Future directions and therapeutic implications

Our findings open several important avenues for future investigation. First, determining the molecular mechanisms underlying circadian regulation of Kir4.1 function—including identification of post-translational modifications, trafficking pathways, and regulatory proteins that differ between DH and VH—could reveal novel targets for modulating astrocytic K^+^ buffering. Second, extending these studies to in vivo preparations using genetically encoded K^+^ sensors and astrocyte-specific manipulations would clarify how the slice-based findings translate to intact circuits and behavior. Third, investigating whether similar regional and temporal heterogeneity exists in human hippocampus and whether these properties are altered in epilepsy could open the way to novel therapeutic avenues.

From a therapeutic perspective, our findings suggest that strategies to enhance astrocytic K^+^ buffering might need to be both regionally and temporally targeted. Global upregulation of Kir4.1 expression throughout the brain could have unintended consequences in regions where current expression levels are functionally appropriate. Similarly, chronotherapeutic approaches that time interventions to periods of maximum vulnerability—when regional differences in buffering capacity are most pronounced—may prove more effective than continuous treatment.

Gene therapy approaches to selectively enhance Kir4.1 expression in VH astrocytes represent a particularly promising avenue. Adeno-associated viral vectors with astrocyte-specific promoters (Tyurikova et al., 2025) could deliver Kir4.1 cDNA to VH, potentially normalizing its K^+^ buffering capacity to DH-like levels and reducing seizure susceptibility. Alternatively, small molecules that enhance Kir4.1 channel open probability or trafficking to the membrane could achieve similar therapeutic effects through pharmacological means.

## Conclusion

This study establishes that astrocytic K^+^ buffering in the hippocampus is neither spatially uniform nor temporally static but instead exhibits regional and circadian heterogeneity. Ventral hippocampal astrocytes possess intrinsically reduced Kir4.1-mediated K^+^ conductance, resulting in rapid K^+^ accumulation during neuronal activity that is partially compensated by putative enhanced Na^+^/K^+^-ATPase activity and expanded gap junction coupling. These regional specializations do not simply mirror neuronal excitability patterns but may actively shape them, creating distinct computational regimes in dorsal versus ventral hippocampus. Superimposed on this stable regional identity is a circadian layer of regulation that modulates—but does not abolish—regional differences, with maximal divergence during early light phase.

Our findings support the view that astrocytes as active, heterogeneous participants in circuit function whose homeostatic properties are dynamically tuned in space and time. This has implications extending beyond the hippocampus to other brain regions and beyond K^+^ buffering to other astrocytic functions. By revealing how regional and temporal factors converge to determine astrocytic K^+^ regulation, this work provides a mechanistic framework for understanding hippocampal circuit dynamics in health and offers potentially new therapeutic strategies for epilepsy based on targeted modulation of astrocytic function at specific times and locations.

## CONFLICT OF INTEREST

The authors declare that they have no conflict of interest.

## DATA AVAILABILITY STATEMENT

The data that support the findings of this study are available from the corresponding author upon reasonable request.

## Acknowledgements

This work was supported by the European Union’s Horizon 2020 Research and Innovation Program under Grant Agreement Number 956325, the Fondation pour la Recherche Médicale (FRM) FDT202404018111, the Croatian Science Foundation under project number IP-2022-10-8493, and Petroleum Technology Development Fund (PTDF/ED/OSS/PHD/KA/2015/22).

## Notes

### Competing Interest Statement

The authors have declared no competing interest.

### Summary of Updates

The title has changed. Figure 1 was updated with more relevant parameters. Figure 4 was completed. The structure of the result and discussion section was changed to reflect the flow of the story that the authors wanted to tell. References were added.

